# Dawn of segmented bilaterians unveiled by soft-bodied fossils in microbial pseudomorphs from the Basal Cambrian

**DOI:** 10.64898/2026.06.18.733126

**Authors:** Xiaoguang Yang, Deng Wang, Farid Saleh, Zhiliang Zhang, Jie Sun, Wenjing Hao, Kentaro Uesugi, Tsuyoshi Komiya, Xing Wang, Jian Han

## Abstract

It is estimated that the evolution of segmented bilaterians occurred during the Ediacaran, whereas most of their unambiguous body fossils did not appear until Cambrian Stage 3. This fossil gap hampers our understanding of the early history of body segmentation, which is a crucial evolutionary innovation for bilaterians. Trace fossils from the late Ediacaran and basal Cambrian suggest this gap is likely to represent a taphonomic bias, implying that the progenitors of segmented bilaterians that existed within this temporal span were hardly preserved. Here, we report a variety of segmented bilaterians from the lowermost Cambrian of South China. These fossils are preserved as microbial pseudomorphs, rather than as ordinary phosphatization. These findings demonstrate that microbial pseudomorphs represent a major pathway for the preservation of segmented bilaterians during this period, as well as an effective mechanism for overcoming the taphonomic bias that affects micro-animals with delicate, non-biomineralized bodies. Moreover, the newly described animals, among the earliest segmented bilaterians, reveal a high diversity of segmented bilaterians during Ediacaran-Cambrian transition, shedding new light on the evolution of body segmentation.

## Introduction

Segmented bilaterians, i.e., vertebrates, panarthropods, and annelids, are the most successful and widespread groups on Earth since the Cambrian Period. Their origin and early evolution are hotly discussed topics in palaeontological research (*Steiner et al., 2004; Peterson et al., 2008; Chen et al., 2018, 2019; Xian et al., 2026*). As molecular clocks studies suggested, the segmented bilaterians might have originated in the early Ediacaran period (*Cunningham et al. 2017; Howard et al. 2022*). During the Ediacaran period, some of the animal fossils, such as *Spriggia, Parvancorina, Kimberella* and *Yilingia* might represent the early candidates of segmented bilaterians. However, the identity of these organisms is controversial (*Chen et al., 2018; Mángano and Buatois, 2020; Shu and Han, 2020; Runnegar, 2022; Dhungana, 2024*). Trace fossils from the Ediacaran–Cambrian transition (ECT) (*Chen et al., 2018, Mangano and Buatois, 2020*) provide insights on the early behaviors of possible segmented bilaterians. While after Cambrian Stage 3, typical segmented bilaterians, such as panarthropods (*Budd, 1998; Yang et al., 2015; Zhang et al., 2016; Caron and Aria, 2017; Ortega-Hernández et al., 2017; Vannier and Martin, 2017; Ou and Mayer, 2018*) and annelids (*Han et al., 2019; Parry and Caron, 2019; Chen et al., 2020; Nanglu and Caron, 2021*) have already become highly derived, serving as the dominant taxa within marine ecosystems. Thus, it is clear to suggest that the gap between the estimated origin of segmented bilaterians and their great radiation after the Terreneuvian is a result of a taphonomic bias, possibly due to the soft and fragile nature of their ancestors (*Anderson et al., 2023*). Although Konservat-Lagerstätten of the early Cambrian are renowned for exceptional preservation of soft organisms (*Liu et al., 2014; Han et al., 2017; Wang et al., 2025*),the preservation window for primitive segmented bilaterians appears to be both restricted and unfavorable under typical phosphatization pathways within these fossil assemblages, resulting in a prolonged absence of their body fossil record. It was only recently that a study provided the first evidence for segmented bilaterian body fossils appeared in the Early Cambrian (*Xian et al., 2026*).

Herein, we present a variety of the earliest segmented bilaterians body fossils from the basal Cambrian of South China (Figure 1, Figure 1—figure supplement 1, 2). These fossils are millimeter-scale with three-dimensional preserved bodies, demonstrating distinct features of segmented bilaterians, including clear bilateral symmetry, dorsoventral and anteroposterior differentiation, and segmented bodies with paired appendages. The morphology of these animals is distinct from that of the newest described annelid fossils (*Xian et al., 2026*). Moreover, detailed observations and Micro-CT analysis show that all animal fossils appeared as compact aggregations of micron-scale phosphatized granules in particular shapes and size, indicating they are preserved as microbial pseudomorphs, rather than direct phosphatization.

**Figure 1.**
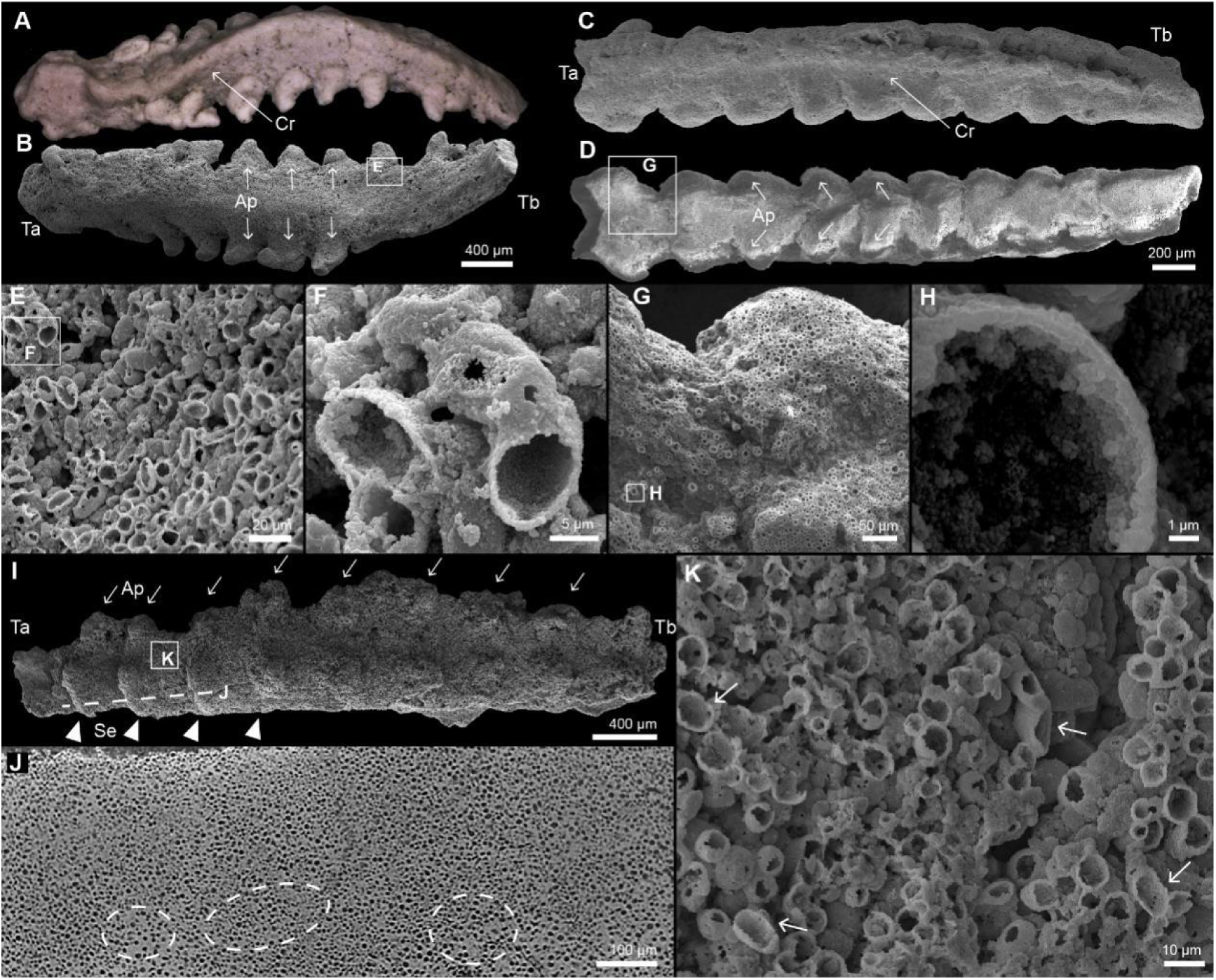
Segmented bilaterians preserved as microbial pseudomorphs from the Cambrian Kuanchuanpu Formation, South China. (A) Dorsal view of ELIXX88-1, with a blunt terminus-A (Ta), a gradually tapered terminus-b (Tb) and a central ridge (Cr). (B) Ventral view of ELIXX88-1, showing nine pairs of appendages. Boxed area enlarged in (E). (C) Dorsal view of ELIXX92-17, with a blunter terminus-a (Ta), a slightly tapered terminus-b (Tb) and a central ridge (Cr). (D) Ventral view of ELIXX92-17, showing nine pairs of appendages. Boxed area enlarged in (G). (E) Compact aggregation of micrograins. Boxed area enlarged in (F). (F) Detailed structure of an ellipsoidal micrograin as a thin-walled vesicle built by phosphate nanocrystals. (G) Compact aggregation of spherical micrograins. Boxed area enlarged in (H). (H) Detailed structure of a spherical micrograin. (I) Ventral view of ELIXX96-21, showing eight recognizable pairs of short cylindrical appendages (Ap) and four recognizable segments (Se), and the biased location of the appendage pair closing to terminus-a within each segment. (J) Tomographic view of the area marked by the dotted line in (I), showing that original animal’s internal structures were completely replaced by micro-granular structures. Dotted line circles mark the solid cement between some granules. (K) Boxed area in (I), showing the mixture of the spherical and ellipsoidal micrograins (arrows) with a domination of spherical type.

Microbes and bacterial films are often associated with the exceptional preservation of soft-bodied fossils through phosphatization (*Wilby et al., 1996; Wilby and Briggs, 1997; Briggs, 2003; Bottjer et al., 2020; Janssen et al., 2022*). And in particular, when microbes, rather than the host’s own tissues, became the main sites of phosphate formation, they created a distinctive “microbial microfabric”. This fabric replaces the original tissues, such as the muscle fibers of arthropods and cephalopods, with dense aggregations of granular bacterial cells (pseudomorph), while still preserving the overall shapes of host organisms (*Wilby and Briggs, 1997; Allison, 1988*). Such pseudomorphing phenomenon has been observed not only in fossils but also under controlled laboratory conditions. Experiments on marine embryos have shown that bacterial biofilms can rapidly assemble and consume them, forming pseudomorphs that replicate detailed embryo morphologies, including cellular structures (*Raff et al., 2008, 2013; Eagan et al., 2017*). Microbial pseudomorph played a significant role in the exceptional preservation of embryo fossils during the ECT (*Raff et al., 2008; Bottjer et al., 2020*), while taphonomic experiments on brine shrimps highlighted their potential for preserving the body structures of micro-animals (*Raff et al., 2013; Eagan et al., 2017*), such as haemocoel and limbs (*Bulter et al., 2015*). The “tightly packed pores” structure and homogeneous internal texture of the latest reported annelid fossils (*Xian et al., 2026*) indicate they are also preserved as microbial pseudomorphs, rather than as endocasts as described in the source literature.

The animal fossils described here, together with the most recently reported annelid fossils (*Xian et al., 2026*), demonstrate that microbial pseudomorphs served as a unique and essential pathway for the preservation of primitive segmented bilaterians with non-biomineralized bodies during the ECT. Furthermore, they help fill a critical gap in our understanding of the evolution of these organisms, revealing a lost world of diversified segmented bilaterians at the dawn of the Cambrian and providing new insights into their early evolutionary history.

## Results

The microfossils described herein were collected from the Fortunian Kuanchuanpu Formation at Zhangjiagou Section, southern Shaanxi Province, China (Figure 1—figure supplement 3), with an estimated age of approximately 535 Ma (*Sawaki et al., 2008*). Among all possible segmented bilaterian fossils (Figure 1; Figure 1—figure supplement 1, 2), the three most representative specimens were selected:

ELIXX88-1: An elongated vermiform body, with a blunt terminus-a (Ta) and a gradually tapered terminus-b (Tb), marking an anterior-posterior differentiation. A central ridge (Cr) occurs along the anterior-posterior (A-P) axis, representing the dorsal side of the body (Figure 1A). Nine pairs of recognizable coniform appendages (Ap) extend laterally to the body (Figure 1B). The terminal parts of appendages bend towards the ventral side with a uniform degree (Figure 2A, 2B, 2G). Therefore, three orthogonal body axes of this specimen could be recognized, suggesting its bilaterian affinity (*Nielsen, 2012*).30 The uniform shape and gesture of the appendages indicate they are repetitive, implying a segmented trunk. The flexure of the main body and appendages shows that this individual was pliable when alive.

**Figure 2.**
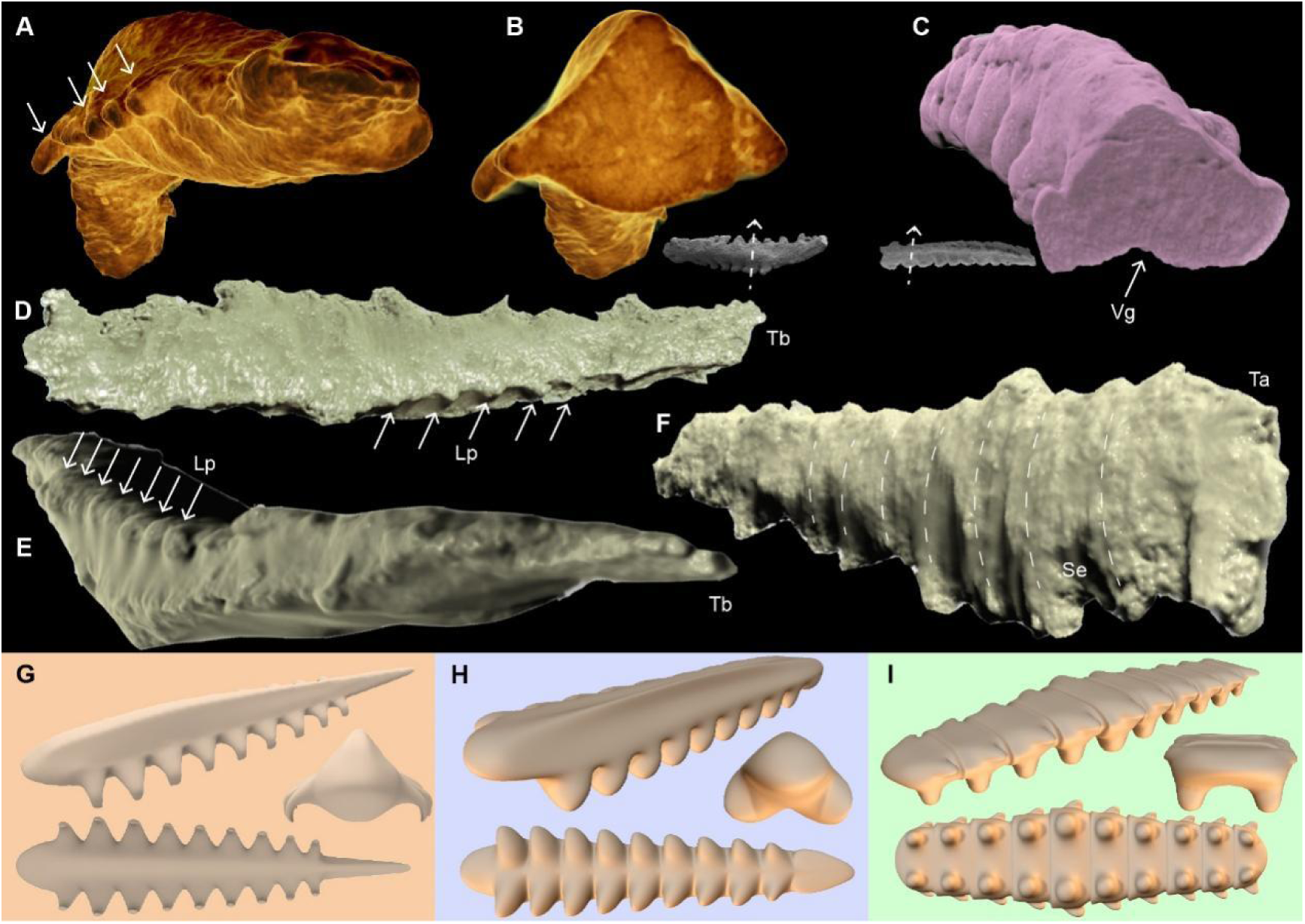
Three-dimensional models based on CT data of segmented bilaterians from the Cambrian Kuanchuanpu Formation, South China. (A) Lateral-front view of ELIXX88-1, showing the appendages located at the ventro-lateral side and extending laterally, with terminal parts bending towards the ventral side. (B) Cross-section of ELIXX88-1, demonstrating the bilateral symmetry of its body and appendages. (C) Transverse section of ELIXX92-17, showing appendages extending semi-laterally from the ventral side and the ventral groove (Vg). (F-H) Dorsal and lateral views of ELIXX96-21, showing its body segments (Se) and lateral protrusions (Lp). (G-I) Reconstruction models of ELIXX88-1, ELIXX92-17, and ELIXX96-21, with overall view (upper), front view (middle), and ventral view (bottom).

ELIXX92-17: A rod-like body with a blunt (possibly cracked) terminus-a and a slightly tapered terminus-b. A central ridge runs the length of the body and indicates the dorsal side (Figure 1C). Nine remaining pairs of clove-shaped appendages extend semi-laterally from the ventral side, with each terminal part pointing towards terminus-a, forming a ventral groove (Vg) along the A-P axis (Figure 1D; 2C and 2H). Although the size of the appendages varies gradually, their shape and orientation remain almost uniform, presumably representing repetitive paired limbs correlated with the body segmentation. The slightly curved body also implies the soft nature of the original animal.

ELIXX96-21: An elongated and flattened body with 10 recognizable segmentations (Se) (Figure 1I). The size of each segment gradually decreases from the central body to both termini. Within each segment, a pair of short cylindrical appendages extends vertically, marking the ventral side. The biased position of each pair of appendages closing to one side of the segment can indicate an anterior-posterior differentiation of the main body (Figure 1I; 2I). A series of conical lateral projections (Lp) is present on the body’s lateral surface close to the dorsal side. Body segmentations are marked by transverse grooves and ridges (Figure 2D-F).

All specimens are comprised of compact aggregations of micron-scale granule structures with hollow interiors (Figure 1E-H,1J, 1K; Figure 1—figure supplement 1E; Figure 1—figure supplement 2G-L). Each granule has a clear boundary and neither intersects nor penetrates other granules. Whether on the surface or in the interior, the granules that constitute each fossil make only tangential contact with one another (Figure 1E, 1G, 1K), or are slightly discrete with solid cement (Figure 1J; Figure 1—figure supplement 1E), thus forming homogeneous bulks. The observed fabrics are comparable to microbial biofilms with fossilized bacteria (granules) and cement of extracellular polymeric substances (EPS) (*Raff et al., 2008*).The granules are thin-walled, rich in both calcium and phosphorus (Figure 1—figure supplement 4F and G), manifesting in phosphate nanocrystals, and have two separate morphogroups, termed “Type-A” and “Type-B”: Type-A granules are ellipsoidal, averaging approximately 14 μm along their long axis and 7 μm along their short axis (Figure 1E, 1F, Figure 1—figure supplement 4D and H); Type-B granules are spherical with an average diameter of ∼9 μm (Figure 1G, 1H; Figure 1—figure supplement 4B and H). Most specimens comprise a single granule type. Rarely, some Type-A granules occur within masses of Type-B (Figure 1K, Figure 1—figure supplement 2J).

## Discussion

### Mechanism for tissue replication in microbial pseudomorphs

The observed granules in the investigated fossils exhibit uniform morphologies, specific size ranges, and hollow interiors (Figure 1, Figure 1—figure supplement 1, 2, 4; Table supplement 1), which are key characteristics of fossilized bacterial cells (*Barling et al., 2023*).These granules can be distinguished from biomorphic micro-grains generated by abiotic phosphatization, which typically display wide and random size distributions, needle-like radiating crystallites, intersecting framboids, and irregular or scattered aggregations (*Baturin and Titov, 2006; Schiffbauer et al., 2012; Cosmidis et al., 2013*). Therefore, these granular microstructures are phosphatized microbe cells rather than diagenetic artifacts.

The highlight of Kucanchuanpu microbial pseudomorphs is that they illustrate a case in which bacterial pseudomorphs replicate almost the entire micro-animal bodies.Despite pristine preservation of external morphology in the investigated fossils as pseudomorphs, their internal anatomy is not preserved. The internal anatomies of these individuals were likely consumed by coccoid (spherical granules) or bacilliform (elongated/ellipsoid granules) bacteria. Following the death of the animal and the rupture of the gut wall, bacteria from the digestive tract proliferated and consumed most internal organs inside out (*Raff et al., 2013; Butler et al., 2015; Eagan et al., 2017; Saleh et al., 2021*). This process forms a centrifugal decay pattern (starting from the inside out), which resulted in a complete loss of internal morphology (*Saleh et al., 2020*). This interpretation is supported by the observation in one specimen of dense phosphate minerals precipitated in its central region (Figure 3F-H), suggesting that the biofilm had the most time to develop at the animal’s core before expanding outward through the body cavity (*Saleh et al., 2020*).

**Figure 3.**
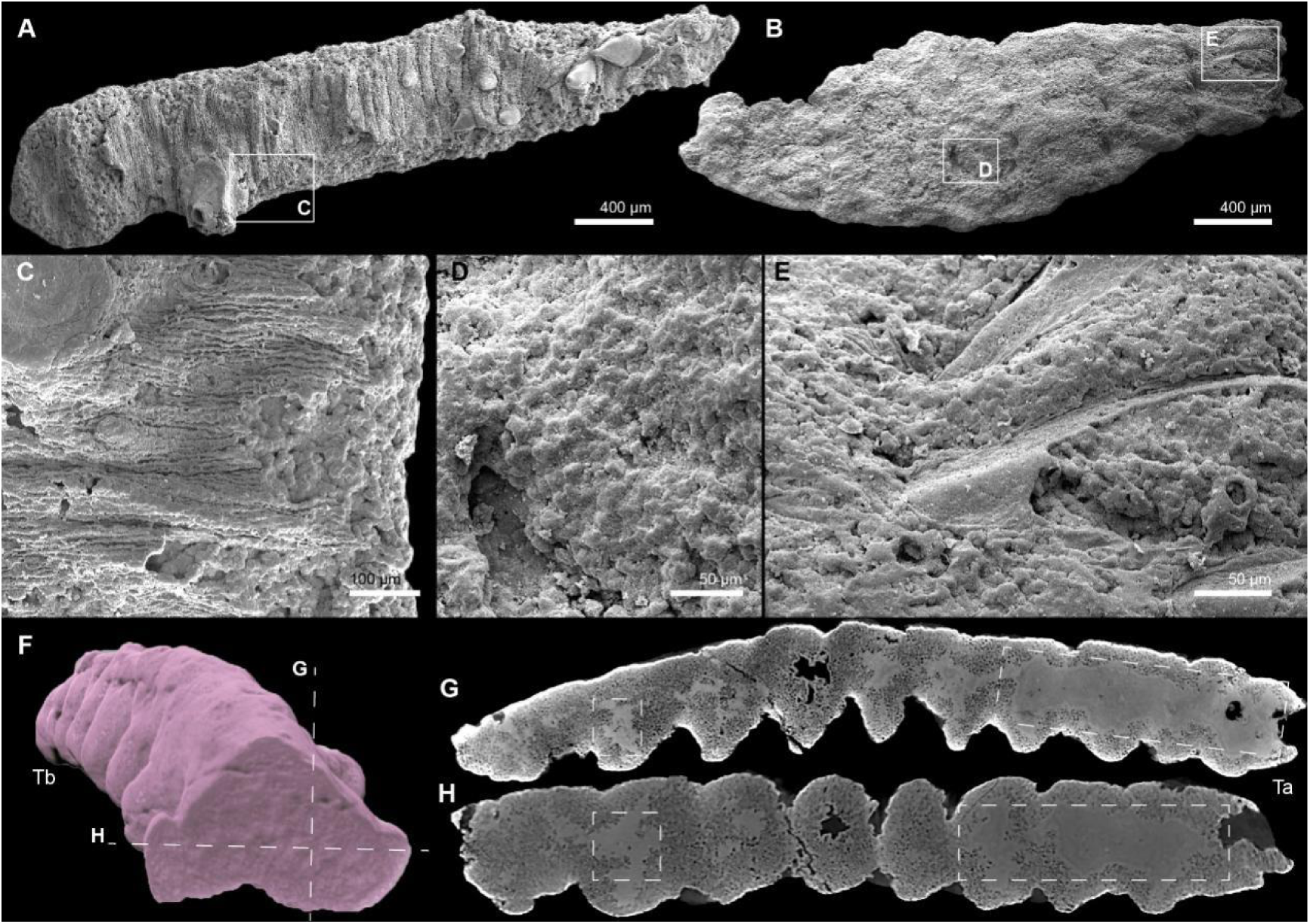
Supporting information on the pseudomorphing mechanism of soft-bodied animals. (A) Part of a scalidophoran worm with dilapidated cuticle and distinct sclerites (ELIXX117-4). (B) Part of a scalidophoran worm with imprints of sclerites and sclerite debris (ELIXX131-60). (C and D) Micrograins that comprise the pseudomorphed worm bodies. (E) Sclerite debris remained on the surface of a pseudomorph. (F) Position indicators of cross sections displayed in (G) and (H) of ELIXX92-17. (G and H) Tomographic views, showing dense bulks of phosphate precipitation in the central part of the animal body.

In such a scenario, a good replication of gross external morphology by bacterial pseudomorph would have benefited from the presence of a more resistant integument, such as the cuticle in a segmented bilaterian. The integument provided structural integrity, allowing microbes to spread into the hemocoel until the entire body was filled with a dense biofilm (*Butler et al., 2015*). This interpretation aligns with the co-occurrences of micro-granular structures and remaining cuticles in some scalidophoran fossils from the same deposit (Figure 3A-E). The integument also likely maintained an enclosed microenvironment under reducing conditions favorable for apatite precipitation (*Corthésy et al., 2025*). In other words, organic matter decay in these organisms was accompanied by biofilm development and phosphate precipitation. Under these conditions, the preservation of the external morphology was likely facilitated by the durable nature of the integument.

Determining the taxonomic affinities of these bacteria involved in this preservation based solely on morphology is challenging despite a growing body of work investigating the role of microbes in exceptional fossil preservation. For instance, modern taphonomic experiments demonstrated that proteobacteria dominate the internal systems of many micro-animals, such as shrimp, and are inefficient at recycling polysaccharides within arthropod integuments (*Corthésy et al., 2024*). Proteobacteria, especially gamma-proteobacteria, are also known as the primary pseudomorph producers (*Eagan et al., 2017; Raff et al., 2008, 2013*). Furthermore, the Gram-negative bacteria, which all proteobacteria belong to, could provide preferential nucleation sites for calcium phosphate by the anionic organic polymers within their periplasm, causing mineralization to start within the bacterial cell walls with hollow encrustations as observed in the investigated fossils (*Fortin et al., 1997; Southam & Donald, 1999; Benzerara et al., 2004; Cosmidis et al., 2013*). Therefore, endogenous proteobacteria are presumed to be the most likely strains responsible for these Cambrian animal pseudomorphs. Although an internal bacterial source has been inferred to be responsible for the formation of pseudomorphs, an external, environmental bacterial source (*Eagan et al., 2017; Raff et al., 2013*) cannot be entirely excluded. Such bacteria may have contributed to the centrifugal decay pattern by first entering the body through openings such as the mouth or the anus.

Interestingly, the observed granules in the investigated fossils are larger than bacteria cells observed in taphonomy experiments (*Wilby & Briggs, 1997; Raff et al., 2008, 2013; Eagan et al., 2017*). One hypothesis to explain this discrepancy is that bacteria, especially proteobacteria, can expand their size due to a fast developmental rate in nutrient-rich environments (*Levin and Angert, 2015; Harris and Theriot, 2018*), such as animal coeloms full of decaying organic material. For instance, *Salmonella* can double in size (*Levin and Angert, 2015*), and in *Escherichia coli*, another typical endogenous gamma-proteobacteria, cell volume can increase over 10 times without changing the morphology (*Si et al., 2017*). The size of the fossil granule might increase further as a result of a mineralization front (where phosphorus and calcium meet to precipitate) that extends beyond the bacterial cell’s immediate boundary (*Raiswell et al., 1993*) (Figure 4C).

**Figure 4.**
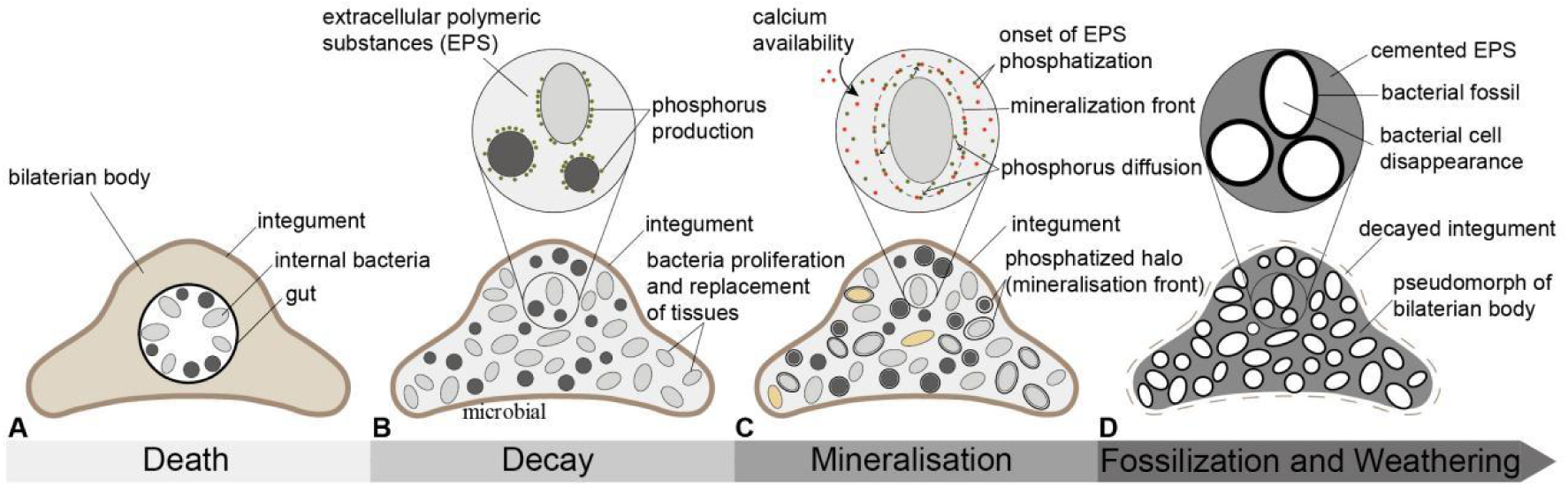
Mechanism of forming an animal body pseudomorph. Following the death of the organism (A), internal bacteria begin to proliferate, releasing phosphorus from the tissues during their decay and secreting extracellular polymeric substances (EPS), which form the biofilm (B). When calcium becomes available, it reacts with the phosphorus, initiating phosphatization (C). Note that if calcium and phosphorus react beyond the immediate boundaries of the bacterial cells, the mineralization front may cause the bacteria to appear larger than they actually are. (D) In the final stages of preservation and weathering, phosphatization continues in both the bacteria and the EPS, while the remaining organic material from the integument is washed away.

In summary, endogenous proteobacteria, possibly aided by some exogenous bacterial taxa, began to degrade the internal organs of a bilaterian animal after death, forming dense biofilms and releasing phosphorus from the tissues into the surrounding environment (Figure 4A, 4B). Simultaneously, the integument remained largely unaffected. Then, the fast-growing microbial colonies crammed against the integument and replicated the animal’s profile (Figure 4B). When conditions became favorable for phosphatization, for instance, with the influx of calcium, minerals began precipitating, and the extracellular polymeric substances were also mineralized, cementing all bacterial cells together (Figure 4C, 4D). The integument was eventually lost with time, either due to decomposition by other microorganisms (*Raff et al., 2013; Corthésy et al., 2024*) or due to alteration over geological times (*Purnell et al., 2018; Saleh et al., 2021*). While the formed pseudomorph persisted due to its extraordinary stability against those unfavorable conditions (*Raff et al., 2013*). This eventually led to fossils preserved in bacterial pseudomorphs replicating the outlines of the original animal body but missing information on internal morphologies (Figure 4D).

### Implications for the early evolution of segmented bilaterians

The newly described Kuanchuanpu micro animals retain key features of their original body plan, such as bilateral symmetry, body differentiation, and serial, repetitive units. All of these characters provide univocal evidence to assign these fossils to segmented bilaterians, which are characterized by serial repetitive organs (*e.g.*, body wall, musculature, nervous systems), tissues, cell types, or body cavities (*Budd, 2001; Blair, 2008*) and typically comprise panarthropods, kinorhynchs, annelids, and chordates. Although the preservation pathway through bacterial pseudomorphs entails the loss of key diagnostic features, including internal anatomy and fine external structures (e.g., possible sclerites, spines, and setae), and thereby hampers very precise phylogenetic placement of the specimens, the fossils, through their preserved gross morphology, nonetheless document the presence of segmented bilaterians with considerable diversity at the onset of the Cambrian (Figure 5, Figure 1—figure supplement 1, 2).

**Figure 5.**
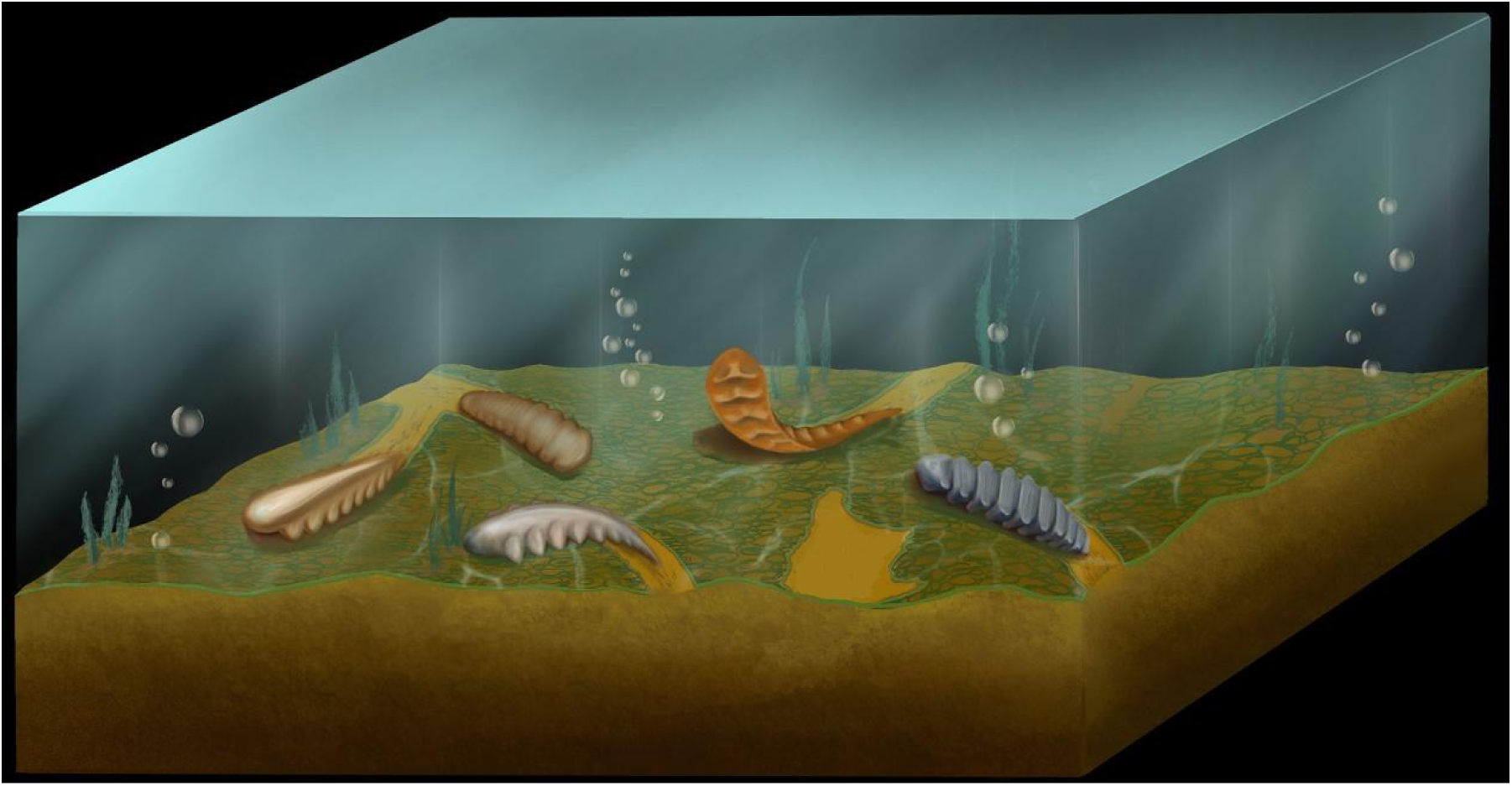
Presumed palaeoecological reconstruction art with the typical early Cambrian segmented bilaterians. From left to right: ELIXX92-17, ELIXX59-71 (in Figure 1—figure supplement 1), ELIXX88-1, ELIXX73-713 (in Figure 1—figure supplement 1), ELIXX96-21.

Molecular clocks estimate the origin of segmented bilaterians to the early Ediacaran (*Cunningham et al., 2017; Howard et al., 2022*). However, the reported early Ediacaran fossil cases of possible segmented bilaterians are controversial. For example, the Ediacaran *Dickinsonia*, bears clear offset segments but glide symmetry. *Kimberella* exhibits bilateral symmetry but only possible internal metameric lobes (*Fedonkin et al., 2007*). *Yilingia* shows a possible modular body (*Dunn et al., 2019 versus Chen et al., 2019*), which contrasts with typical later bilaterians in the Cambrian (*Shu and Han, 2020*), and its segmentation and appendages might need more assessment (*Mangano and Buatois, 2020*). More importantly, none of the aforementioned examples have all the fundamental features of true segmented bilaterians (i.e., bilaterality and serial, repetitive units) at the same time.

The diversity in the bilaterian material presented herein helps in filling the gap between molecular data and the fossil record, by indicating that the evolution of body segmentation commenced during the Ediacaran period while segmented bilaterians became diversifying at the ECT. This is also corroborated by trace fossil data. Trace fossils with bilobate and paired grooves in the basal Cambrian, such as *Rusophycus* and *Cruziana* (*Mángano and Buatois, 2020; Holmes and Budd, 2022*), demonstrate that segmentation had already evolved in some different types of bilaterians, regardless of the affinities of their tracemakers (e.g. arthropods or annelids) (*Donovan et al., 2010; Knaust, 2022*).

Furthermore, our specimens exhibit features of homonomous segments. The components of each body segment (i.e. appendages) repeated simply, without change in structures. These fossils likely indicate a transformation from an incipient evolutionary stage toward more derived heteronomous segmentation, such as arthropods and some annelids during the peak of the Cambrian explosion. This suggests that segments of early segmented animals possess evolutionary plasticity (*Deline et al. 2018; Peterson et al. 2008; Chipman 2010*) and segmented bilaterians that dominated marine ecosystems from the Cambrian Stage 3 might have evolved from a simple segmental body by a stepwise (i.e., “system by system”) evolution (*Budd, 2001; Chipman, 2019*).

## Materials and Methods

All specimens were recovered from the Bed 2 of phosphorus limestone of the Kuanchuanpu Formation at the Zhangjiagou Section, Dahe Town, Xixiang County, Hanzhong City, Shaanxi Province, China (Figure 1—figure supplement 1). The Xixiang area was palaeogeographically located on the northwestern margin of the Yangtze Platform during the Ediacaran and Cambrian periods and the fossil-sampling horizon can be correlated with the Anabarites trisulcatus-Protohertzina anabarica assemblage zone, which indicates the Fortunian Stage with an estimated age of 536.5 ± 2.5 Ma (*Sawaki et al. 2008*). Rock samples were treated with 8-10% acetic acid solution and phosphatized fossils were then selected from the residues using a binocular microscope. All specimens were deposited in the collections of the Early Life Institute of Northwest University, Xi’an, China.

### Scanning electron microscopy (SEM)

Selected fossils were coated with gold then observed with an FEI Quanta 450 FEG Scanning Electron Microscope (SE Mode, high vacuum, 20 kV, spot size 4.0) at State Key Laboratory of Continental Dynamics, Northwest University, Xi’an, China.

### X-ray computed microtomography and 3D reconstruction

High-potential specimens were analyzed with a ZEISS Xradia-520 micro-computed tomography (50 kV/4W, 80 μA, 1.10×1.10×1.10 to 4.23×4.23×4.23 μm^3^ voxel^−1^ depended on fossil size) at the State Key Laboratory of Continental Dynamics (SKLCD), Northwest University, China and one specimen XX59-71 was analyzed at SPring-8 in Hyogo, Japan (23 KeV, 0.49×0.49×0.49 μm^3^ voxel^−1^). X-ray microtomography data were processed through Dragonfly 3.5 software for detailed tomographic analysis and making three-dimensional visualization pictures and videos.

### Measurements

Statistical analysis were based on the micrograins which exhibit relatively intact three-dimensional outlines and diameters could be determined. Measurements for the grain size were performed by ImageJ 1.53K with selected SEM images.

## Data availability

All original data of Micro-CT and SPring-8 analysis that support this study have been deposited in Figshare: (https://figshare.com/s/b1a9aa032d009fdbeaac). All figured specimens are housed in the collections of the Early Life Institute of Northwest University and can be accessed by contacting Jian Han. All data are available in the main text or the supplementary files.

## Acknowledgments

We thank J. Luo (NWU) for assistance with fossil collection, X. Liu (NWU) and W. Zhou (XAUT) for the artistic reconstructions, Profs. J. Paterson (University of New England) and Q. Ou (China University of Geosciences Beijing) for constructive comments during draft writing and editing. This publication was supported by the National Natural Science Foundation of China (grant numbers 41902012, 42202009, 42372012, 41720104002 and 42402006); National Key research and Development Program of China (grant number 2023YFF0803601); High-Level Talents of Liupanshui Normal University (grant numbers LPSSYKYJJ202506).

## Funding

National Natural Science Foundation of China (41902012, 42202009, 42372012, 42402006 and 41720104002)

National Key research and Development Program of China (2023YFF0803601)

High-Level Talents of Liupanshui Normal University (LPSSYKYJJ202506)

## Author contributions

Xiaoguang Yang, Conceptualization, Formal analysis, Investigation, Methodology, Writing-original draft, Writing-review and editing, Funding acquisition; Deng Wang, Investigation, Writing-original draft, Writing-review and editing, Funding acquisition; Farid Saleh, Investigation, Methodology, Writing-original draft, Writing-review and editing; Jie Sun, Formal analysis, Visualization; Wenjing Hao, Formal analysis, Visualization, Writing-review and editing; Kentaro Uesugi, Formal analysis, Methodology; Tsuyoshi Komiya, Formal analysis, Methodology; Xing Wang, Conceptualization, Formal analysis, Investigation, Writing-review and editing, Funding acquisition; Jian Han, Conceptualization, Supervision, Data curation, Writing-review and editing, Funding acquisition.

## Supplementary Files

**Figure 1—figure supplement 1.**
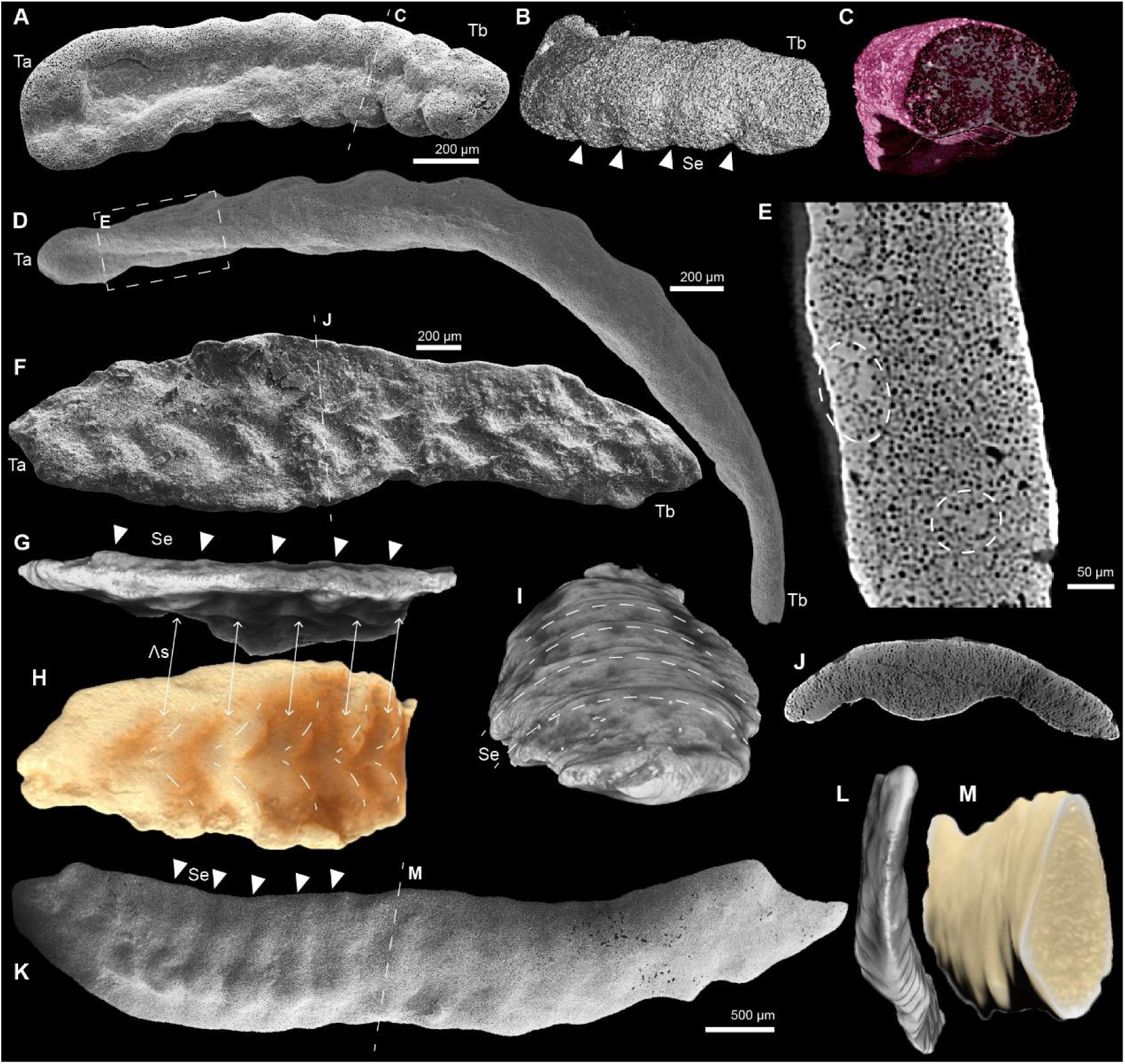
Possible bilaterian fossils with paired appendages or segmented bodies. (A) Ventral view of ELIXX59-71, showing a blunter terminus-a (Ta) and thinner terminus-b (Tb), with a distinct groove along the body axis and five pairs of recognizable stubby appendages. (B) Three-dimensional CT model with dorsal view of ELIXX59-71, showing clear segments (Se) correlating with each appendage pair. (C) Transversal view of the dotted line area in (A). (D) Dorsal view of ELIXX56-367, with a rounded terminus-a (Ta) and a long, gradually tapered terminus-b (Tb). (E) Tomographic view of the area marked by the dotted line box in (D), showing homogeneous aggregation of spherical micrograins and solid cements between micrograins (dotted line circles). (F) Ventral view of ELIXX73-713 with a wider terminus-a (Ta) and a narrower terminus-b (Tb), presenting pairs of “Λ” shaped structures (Λs), possibly related to paired appendages. (G-H) Lateral and ventral views of the three-dimensional CT model of the cracked left part of ELIXX73-713, indicating each structure (Λs) occurred between two segments (Se). (I) Dorsal view of CT model in (G), showing segments (Se) marked by transverse ridges. (J) Tomographic view of the section marked by the dotted line in (F), demonstrating a bilateral body. (K) Lateral view of ELIXX84-262, with a segmented trunk. (L) Three-dimensional CT model of ELIXX84-262, showing the bilateral symmetry of segment distributions. (M) Transversal view of dotted line area in (K), showing the bilateral symmetry of the body.

**Figure 1—figure supplement 2.**
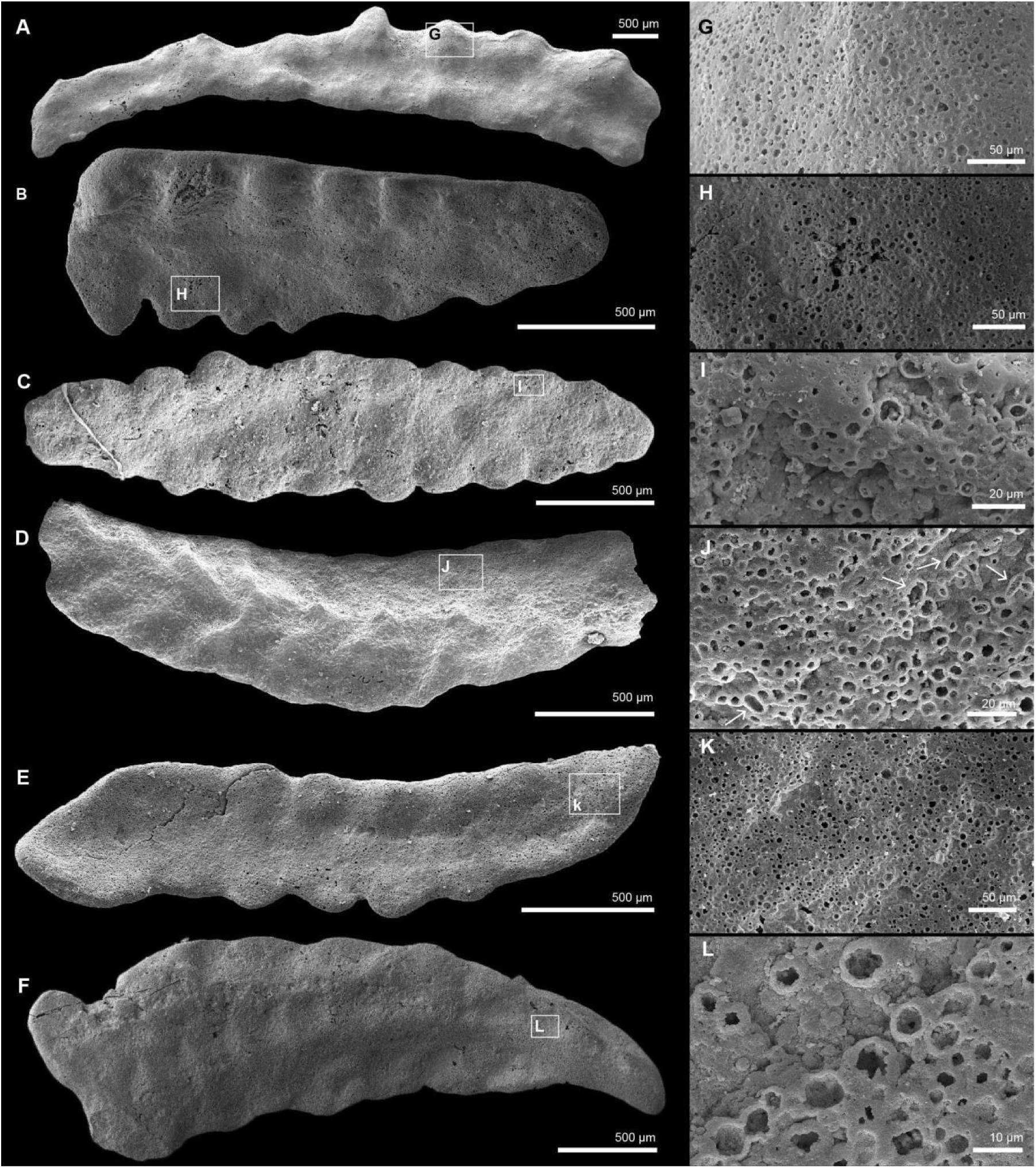
Additional possible segmented bilaterians from the Cambrian Kuanchuanpu Formation, South China. (A-F) ELIXX139-140, ELIXX133-2, ELIXX117-21, ELIXX117-117, ELIXX135-186, ELIXX91-63. (G-L) Zoomed-in views of box areas in each specimen, showing the preservation type as microbial pseudomorphs.

**Figure 1—figure supplement 3.**
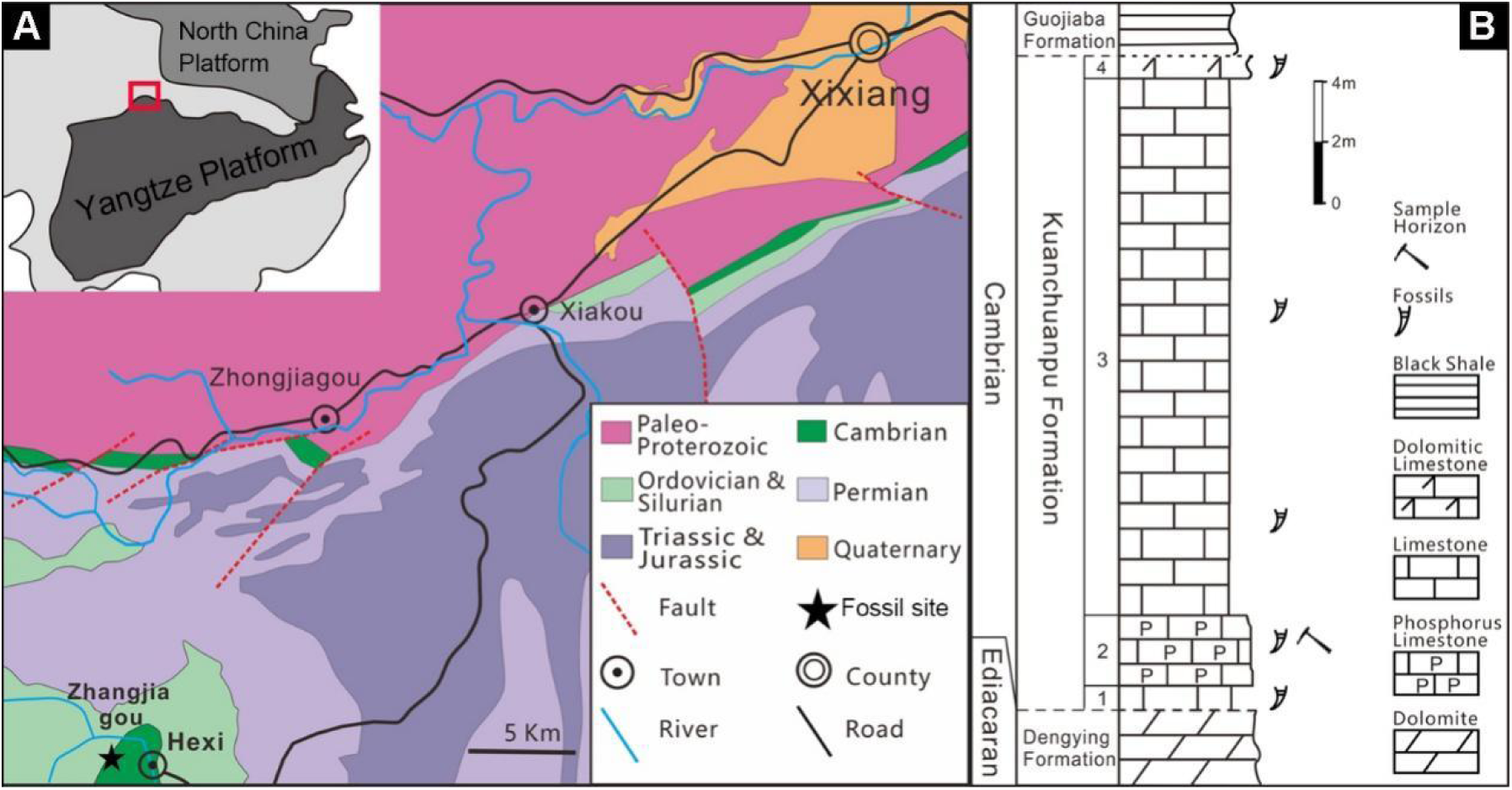
Fossil locality and stratigraphy of the Zhangjiagou section in Xixiang County, Shaanxi Province, China. (A) Palaeogeographical position and geological map of the Zhangjiagou section. (B) Stratigraphic column of the Zhangjiagou section with sampling horizon marked by a hammer icon.

**Figure 1 — figure supplement 4.**
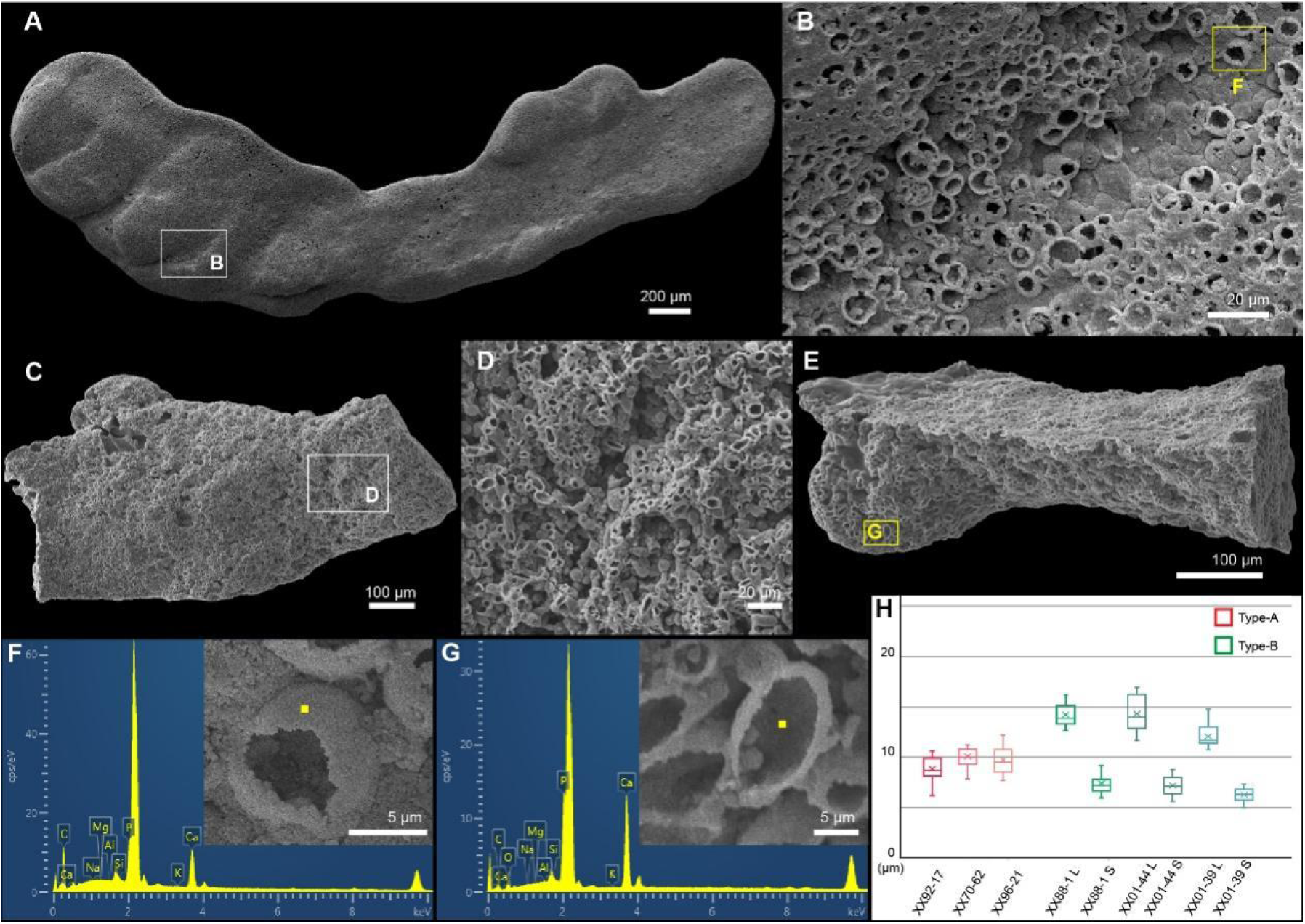
Additional information on the constitutive micrograins of animal pseudomorphs from the Cambrian Kuanchuanpu Formation, South China. (A) Specimen ELIXX70-62, an unrecognizable animal with possible segments. (B) Many three-dimensionally preserved spherical micrograins from the box in (A). (C and E) Specimens ELIXX01-44 and 01-39, Pseudomorph fragments consisted of ellipsoidal micrograins. (D) Enlarged view of the boxed area in (C). (F and G) Energy-dispersive X-ray spectroscopy maps of boxed areas in (B) and (E), showing the major constituent of the micrograins as Ca-phosphate. (H) Statistical results for the size distribution of the spherical and ellipsoidal micrograins. L: long axis, S: short axis.

**Table supplement 1.**
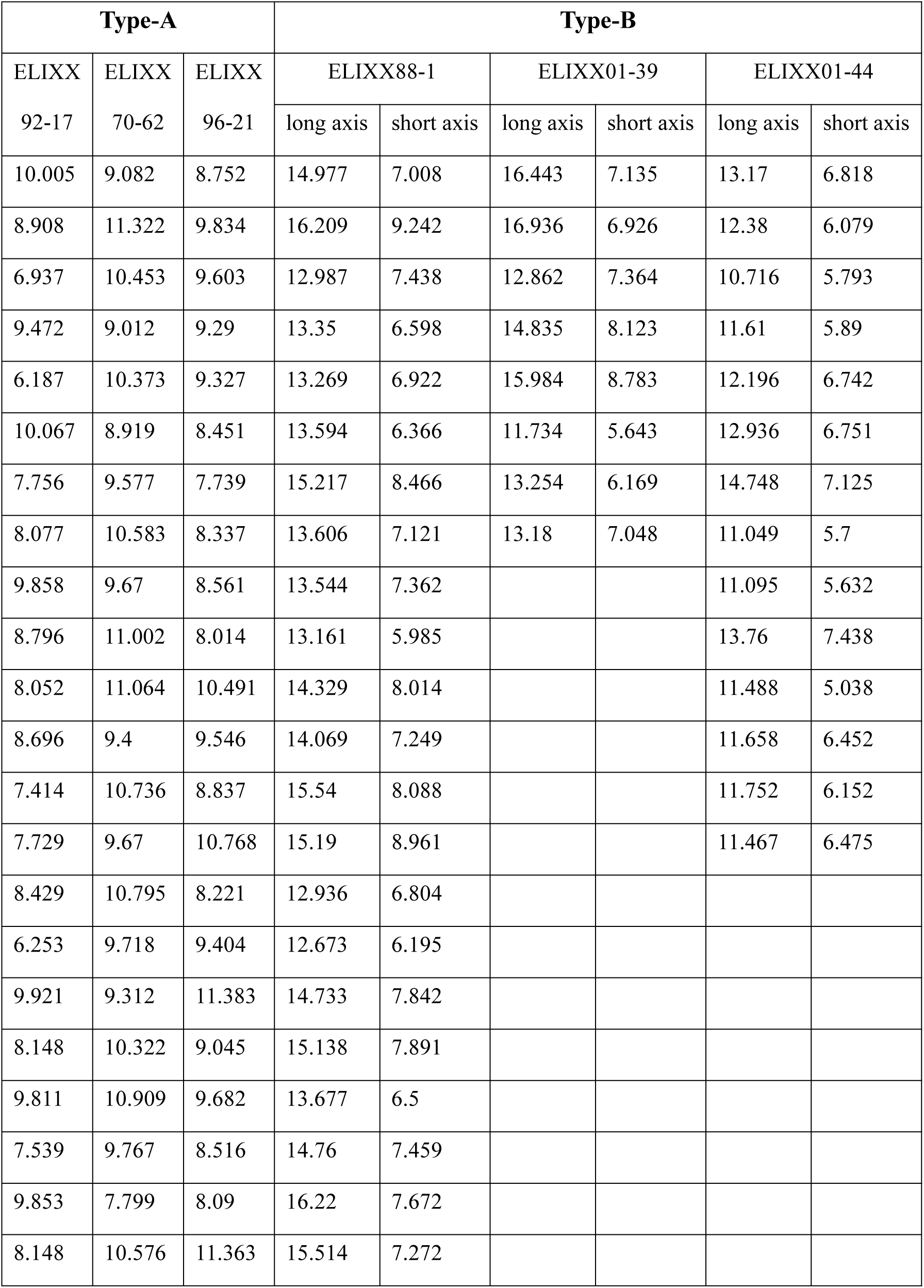
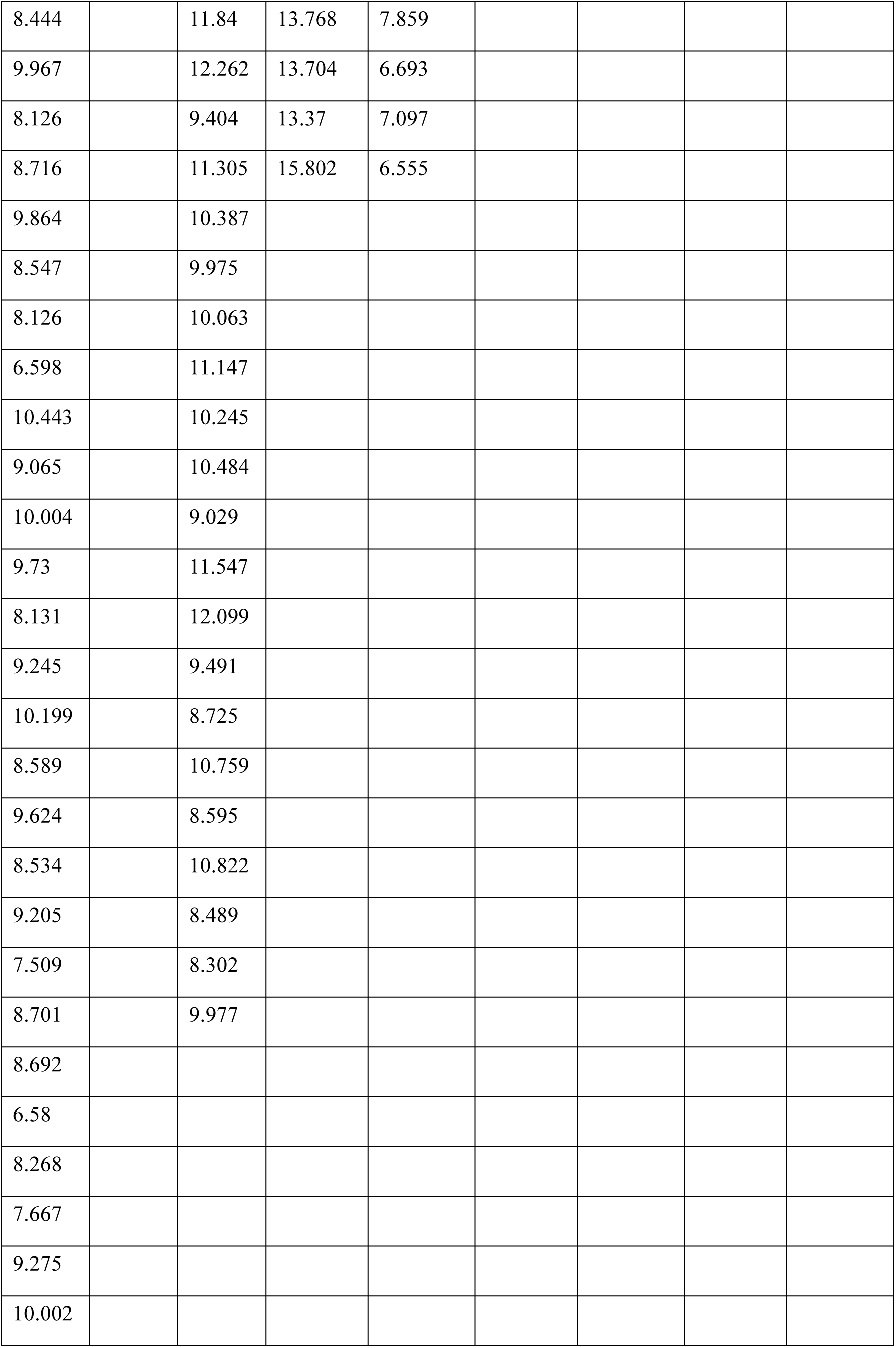
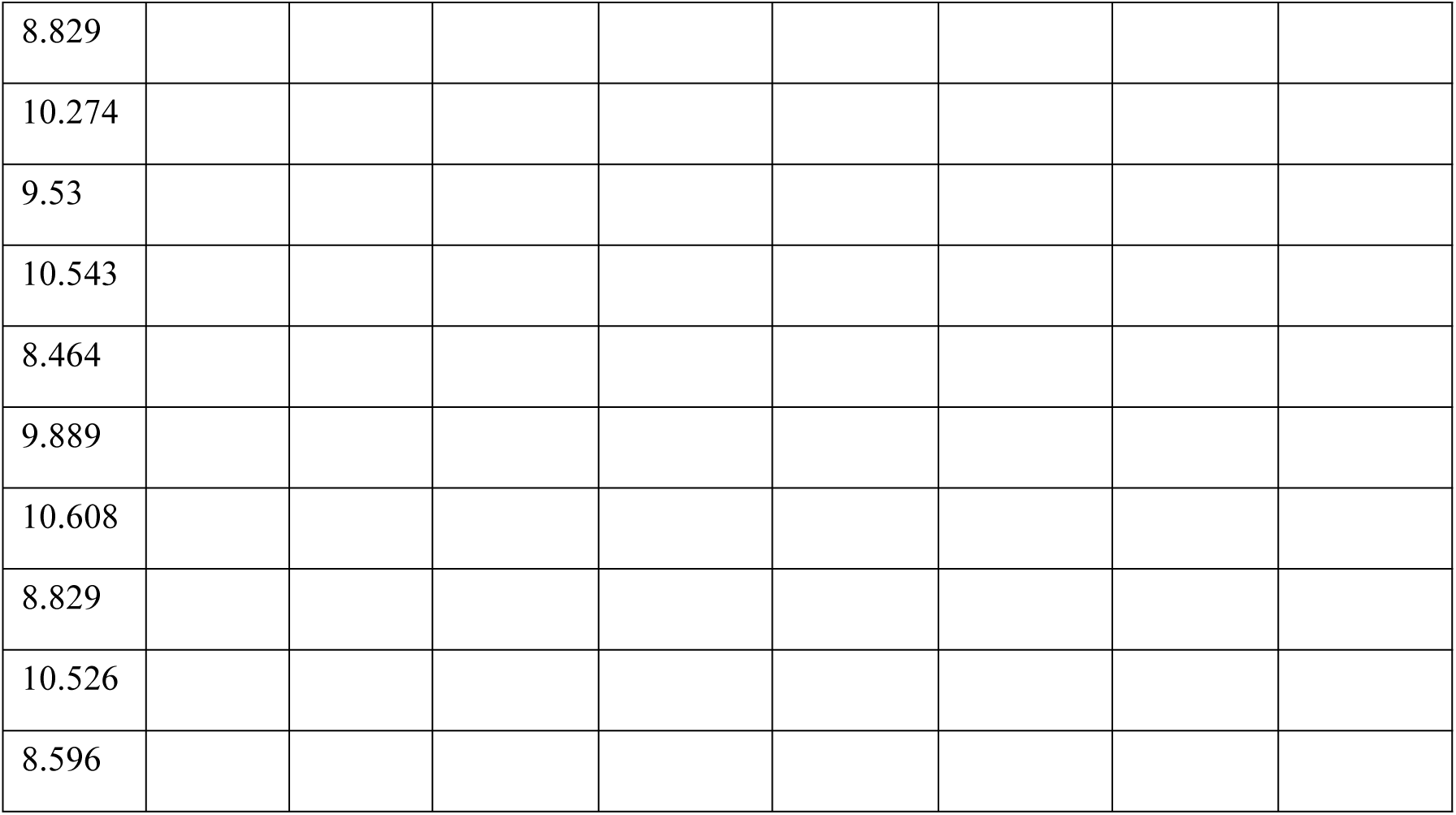
Size of constitutive micrograins of pseudomorphs from the Cambrian Kuanchuanpu Formation, South China (μm)

## Supplementary videos

**Movie S1.**

3D visualization movie of specimen ELIXX92-17.

**Movie S2.**

3D visualization movie of specimen ELIXX88-1.

**Movie S3.**

3D visualization movie of specimen ELIXX96-21.

**Movie S4.**

3D visualization movie of specimen ELIXX59-71.

**Movie S5.**

3D visualization movie of specimen ELIXX56-367.

**Movie S6.**

3D visualization movie of specimen ELIXX73-713.

**Movie S7.**

3D visualization movie of specimen ELIXX84-262.

